# Unraveling Fe(II)-oxidizing mechanisms in a facultative Fe(II) oxidizer *Sideroxydans lithotrophicus* ES-1 via culturing, transcriptomics, and RT-qPCR

**DOI:** 10.1101/2021.08.18.456919

**Authors:** Nanqing Zhou, Jessica L. Keffer, Shawn W. Polson, Clara S. Chan

## Abstract

*Sideroxydans lithotrophicus* ES-1 grows autotrophically either by Fe(II) oxidation or thiosulfate oxidation, in contrast to most other neutrophilic Fe(II)-oxidizing bacteria (FeOB) isolates. This provides a unique opportunity to explore the physiology of a facultative FeOB and constrain the genes specific to Fe(II) oxidation. We compared the growth of *S. lithotrophicus* ES-1 on Fe(II), thiosulfate, and both substrates together. While initial growth rates were similar, thiosulfate-grown cultures had higher yield with or without Fe(II) present, which may give ES-1 an advantage over obligate FeOB. To investigate the Fe(II) and S oxidation pathways, we conducted transcriptomics experiments, validated with RT-qPCR. We explored the long-term gene expression response at different growth phases (over days-week) and expression changes during a short-term switch from thiosulfate to Fe(II) (90 min). The *dsr* and *sox* sulfur oxidation genes were upregulated in thiosulfate cultures. The Fe(II) oxidase gene *cyc2* was among the top expressed genes during both Fe(II) and thiosulfate oxidation, and addition of Fe(II) to thiosulfate-grown cells caused an increase in *cyc2* expression. These results support the role of Cyc2 as the Fe(II) oxidase and suggest that ES-1 maintains readiness to oxidize Fe(II) even in the absence of Fe(II). We used gene expression profiles to further constrain the ES-1 Fe(II) oxidation pathway. Notably, among the most highly upregulated genes during Fe(II) oxidation were genes for alternative complex III, reverse electron transport and carbon fixation. This implies a direct connection between Fe(II) oxidation and carbon fixation, suggesting that CO_2_ is an important electron sink for Fe(II) oxidation.

**Importance:** Neutrophilic FeOB are increasingly observed in various environments, but knowledge of their ecophysiology and Fe(II) oxidation mechanisms is still relatively limited. *Sideroxydans* are widely observed in aquifers, wetlands, and sediments, and genome analysis suggests metabolic flexibility contributes to their success. The type strain ES-1 is unusual amongst neutrophilic FeOB isolates as it can grow on either Fe(II) or a non-Fe(II) substrate, thiosulfate. Almost all our knowledge of neutrophilic Fe(II) oxidation pathways comes from genome analyses, with some work on metatranscriptomes. This study used culture-based experiments to test the genes specific to Fe(II) oxidation in a facultative FeOB and refine our model of the Fe(II) oxidation pathway. We gained insight into how facultative FeOB like ES-1 connect Fe, S, and C biogeochemical cycling in the environment, and suggest a multi-gene indicator would improve understanding of Fe(II) oxidation activity in environments with facultative FeOB.

## 1. Introduction

Neutrophilic Fe(II)-oxidizing bacteria (FeOB) are increasingly found in a wide variety of terrestrial and marine environments (1–4), often in suboxic zones where microaerophilic FeOB can successfully outcompete abiotic Fe(II) oxidation (5, 6). In these environments, FeOB have the potential to affect many elemental cycles, notably carbon cycling as many neutrophilic FeOB are autotrophic and sequester organics in their biominerals (7, 8). The most well-characterized neutrophilic FeOB are the marine *Zetaproteobacteria* and freshwater/terrestrial *Gallionellaceae* (2, 9, 10), though our knowledge of both remains limited partly because there are relatively few isolates, due to challenges in culturing FeOB. Furthermore, most FeOB isolates are obligate Fe(II)-oxidizers, limiting our understanding of Fe(II) oxidation mechanisms and ecophysiology of facultative FeOB.

Within the *Gallionellaceae, Sideroxydans* is a genus of microaerophilic Fe(II)-oxidizers widespread in terrestrial aquatic systems including groundwater, wetlands, creek sediments, and rhizosphere soil (11–16). *Sideroxydans* have been identified from moderately acidic to circumneutral environments, over a wide range of oxygen concentrations (17–20), commonly in environments with both high Fe and S concentrations (21–23). The type strain is *Sideroxydans lithotrophicus* ES-1, isolated from groundwater using FeS and O_2_, with growth over a pH range of 5.5-7.5 (24). ES-1 is able to grow on Fe(II) or thiosulfate, making it unusual amongst neutrophilic FeOB in that it is a facultative Fe(II)-oxidizer. The strain has a fully sequenced, closed genome that encodes multiple putative Fe(II) oxidases (25), suggesting further metabolic flexibility. The metabolic versatility makes ES-1 a good model to study the physiology of a facultative FeOB, compare the expression of different putative Fe(II) oxidase genes, and investigate genes that are specifically involved in Fe(II) oxidation.

The ES-1 genome includes multiple potential pathways for both S and Fe oxidation. The sulfur oxidases encoded in the ES-1 genome are thiosulfate dehydrogenase (Tsd), reverse dissimilatory sulfite reductase (rDsr), and sulfur oxidase (Sox). The two main putative Fe(II) oxidases in ES-1 are Cyc2 and MtoA (25, 26). Cyc2 is the homolog of the Fe(II) oxidase identified in *Acidithiobacillus ferrooxidans* (27) and *Mariprofundus ferrooxydans* (28), and is widely distributed in microaerophilic FeOB (26, 29). MtoA is the homolog of the Fe(III) reductase MtrA in *Shewanella oneidensis* and showed Fe(II) oxidation activity *in vitro* (30). Previous research suggests microaerophilic Fe(II) oxidation occurs extracellularly (31–33), which requires periplasmic electron carriers to transfer electrons from Cyc2 or MtoA to terminal oxidases or Complex I through the reverse electron transfer (RET) chain (8, 31, 33). Genomic analysis of ES-1 has revealed genes that encode different parts of the potential electron transfer pathways, including *cbb*_*3*_ and *bd* cytochrome oxidases as the terminal oxidases, *bc*_*1*_ complex and alternative complex III (ACIII) as Complex III, NADH dehydrogenase (Complex I) and succinate dehydrogenase (Complex II) (25). Moreover, ES-1 contains genes for two forms of RuBisCO, which would enable it to efficiently fix CO_2_ in different environmental conditions (25). However, since no ES-1 transcriptomics or proteomics has been performed, the Fe(II) oxidation pathway has not been verified. In particular, it is still unknown if *cyc2* and *mtoA* are specifically involved in Fe(II) oxidation and the role of ACIII and other cytochromes has been speculative.

In this study, we used physiological, transcriptomic and reverse transcription quantitative PCR (RT-qPCR) approaches to study the metabolism and metabolic pathways of ES-1 using different substrates. Growth was characterized on Fe and S substrates individually, in series (i.e. substrate switching), and together. Transcriptomes of ES-1 grown on Fe(II) (Fe(II)-citrate and FeCl_2_) or thiosulfate were sequenced to look at the long-term (days) and short-term (min-hr) response to different electron donors. Differential gene expression (DGE) analysis was used to constrain genes specific to Fe(II) oxidation. RT-qPCR was used to further quantify the expression of *cyc2* and *mtoA* in different conditions. With these results, we gained insight into the physiology of a metabolically-flexible Fe(II)-oxidizer, clarified the roles of potential Fe(II) oxidase genes, and improved the current Fe(II) oxidation electron transfer pathway model in a facultative microaerophilic FeOB.

## 2. Results

### 2.1 Characterizing ES-1 growth on different substrates

We compared ES-1 growth on thiosulfate or Fe(II) individually and when both substrates were available concurrently. We grew ES-1 cultures with a daily amendment of only FeCl_2_, only thiosulfate, or both FeCl_2_ and thiosulfate at a daily target concentration of 500 μM. Cell growth rates were initially comparable (slightly faster on FeCl_2_ + thiosulfate at day 2), but ultimately, the cell concentration in the thiosulfate-only culture was the highest (Figure 1A). The only thiosulfate oxidation product was tetrathionate (Figure S1), thus in ES-1, Fe(II) oxidation and thiosulfate oxidation are both one-electron reactions. The electron uptake rate at mid-log phase was 3.7 – 3.8 × 10^−8^ μmol e^-^/cell/day in the cultures with single substrates, which was slightly higher than the culture with both substrates (2.9 × 10^−8^ μmol e^-^/cell/day). When both substrates were present, cells consumed Fe(II) and thiosulfate simultaneously, suggesting there is no preference (red lines in Figures 1B and 1C). However, there was a lag in thiosulfate consumption at day 1 (blue line in Figure 1C), perhaps due to the fact that the inoculum was pre-grown on Fe(II). To investigate if thiosulfate-grown cells would show a lag in Fe(II) oxidation when switched back to FeCl_2_, we grew ES-1 on thiosulfate and then spiked in FeCl_2_. In this case, there was no lag, as thiosulfate-grown ES-1 started to oxidize Fe(II) immediately, while the azide-killed control showed negligible Fe(II) oxidation, demonstrating that the Fe(II) oxidation was nearly all biotic (Figure 1D). The lag from Fe(II) oxidation to thiosulfate oxidation (but not the reverse) suggests that ES-1 maintains readiness to oxidize Fe(II) but regulates its ability to oxidize thiosulfate.

**Figure 1.**
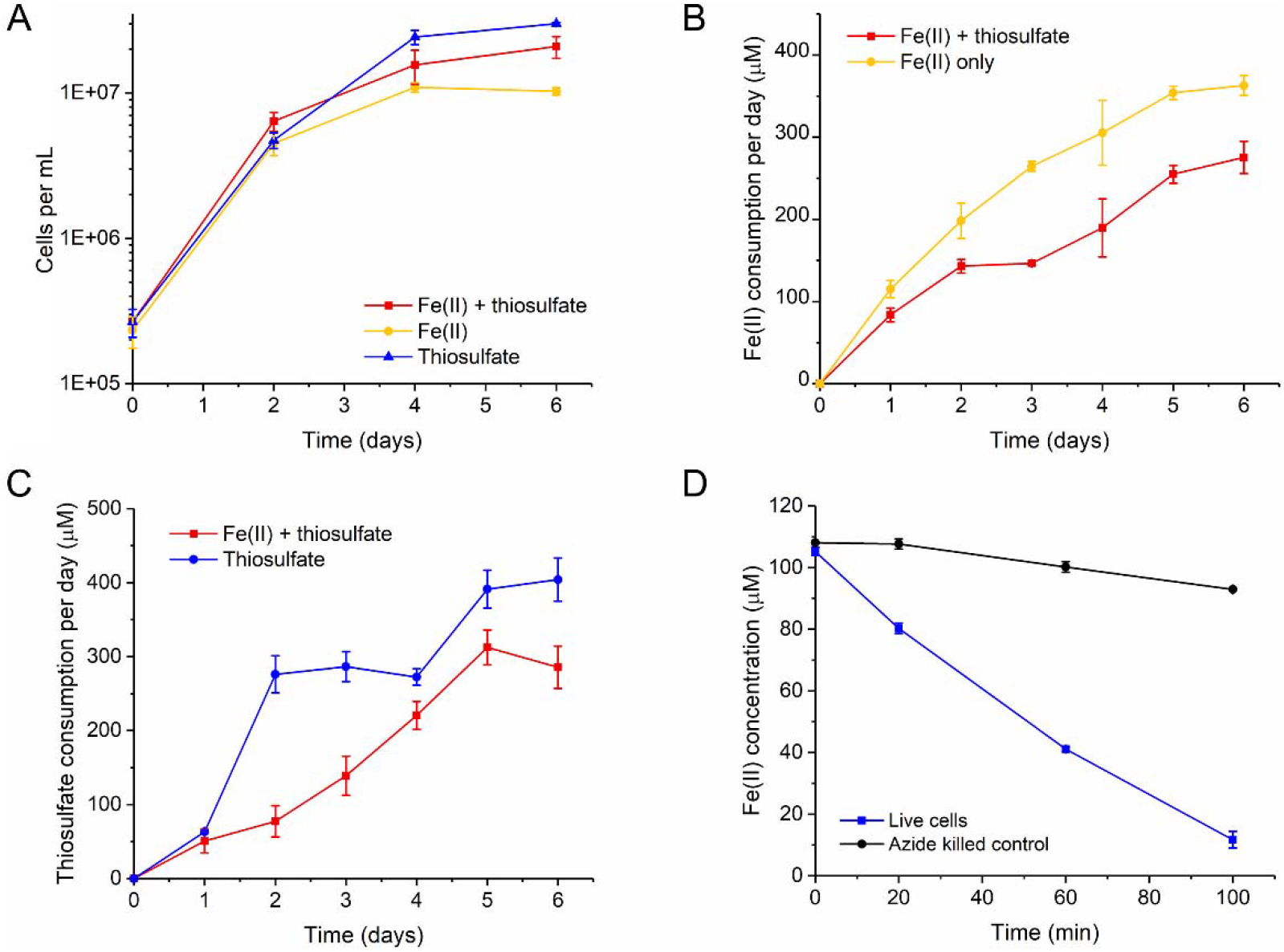
(A) ES-1 growth on Fe(II) only, thiosulfate only, or Fe(II) + thiosulfate culture; (B) biotic Fe(II) consumption per day in Fe(II) only culture and Fe(II) + thiosulfate culture; (C) biotic thiosulfate consumption per day in thiosulfate only culture and Fe(II) + thiosulfate culture; (D) Fe(II) concentration change after spiking FeCl_2_ (100 μM) into stationary phase thiosulfate-grown culture. The Fe(II) and thiosulfate consumption values were calculated using the concentration after FeCl_2_ and thiosulfate supplement on day *n* subtracted by the FeCl_2_ and thiosulfate concentration before supplementation on day *n+1*. To calculate biotic consumption, the abiotic consumption was subtracted from culture consumption values.

Growing ES-1 on FeCl_2_ yields copious oxyhydroxides, which can interfere with RNA extraction. We therefore evaluated the growth of ES-1 on Fe(II)-citrate, as citrate would chelate Fe(III) and prevent mineral formation, but not affect Fe(II) oxidation kinetics (34). ES-1 can grow on Fe(II)-citrate without forming Fe(III) oxyhydroxides but showed no growth on citrate alone, demonstrating that growth was due to Fe(II) oxidation (Figure S2A). We optimized cell growth on Fe(II)-citrate and thiosulfate by trying a series of substrate concentrations (100 μM-750 μM of Fe(II) per day, one time addition of 0.5-10 mM thiosulfate) (Figures S2A and S2B). The growth data showed that when cultures were dosed with similar amounts of Fe(II)-citrate and thiosulfate, yield on thiosulfate was higher. From these results, we chose an Fe(II)-citrate concentration of 500 µM per day and a thiosulfate concentration of 10 mM to maximize the biomass yield on each substrate.

### 2.2 Transcriptome analysis revealed genes used during Fe(II) and thiosulfate oxidation

To investigate genes involved in Fe(II) and thiosulfate oxidation, we conducted long-term single-substrate experiments and a short-term Fe(II) addition experiment. In the long-term experiment (7-10 days), cells were grown on either Fe(II) or thiosulfate, and samples were taken from different growth phases: mid-log, late-log and early stationary phase (Figures 2A and 2B). To analyze differential gene expression, pairs of time-points were compared (for example, mid-log on Fe(II)-citrate to mid-log on thiosulfate). To investigate the short-term response of ES-1 to Fe(II), we spiked FeCl_2_ into an ES-1 culture grown on thiosulfate, and collected samples at the early (15 min), middle (35 min) and the end (90 min) of the Fe(II) oxidation curve, along with a pre-spike time 0 control for comparison (Figure 2C). In this short-term experiment, gene expression at each time-point (15, 35, and 90 min) was compared to that of the thiosulfate-grown stationary cells (time 0). All gene expression values are presented as transcripts per million (TPM), normalized by dividing by the average TPM of six constitutively expressed genes (*gyrB, gyrA, adk, rho, era, gmk*; (35)).

**Figure 2.**
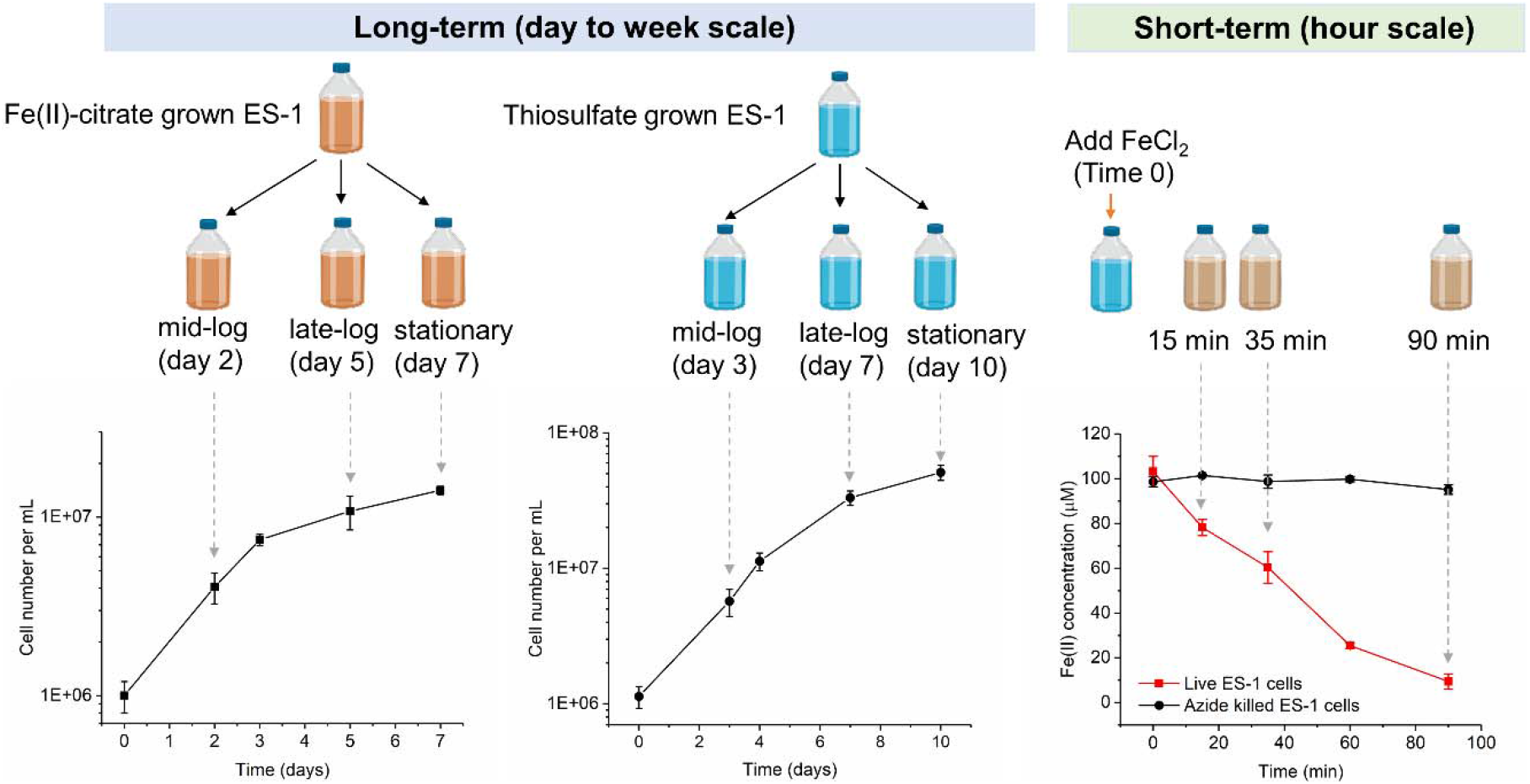
(A) Growth of ES-1 with daily feeding of 500 µM Fe(II)-citrate; (B) Growth of ES-1 on one-time dose of 10 mM thiosulfate; (C) Fe(II) oxidation kinetics after FeCl_2_ spike into stationary thiosulfate culture. Gray arrows indicate the sample points.

Clear differences in gene expression were evident between growth on Fe(II) citrate and thiosulfate in both long- and short-term experiments from the differential gene expression (DGE) analyses. Mid-log phase represented the largest difference between Fe(II)-citrate and thiosulfate cultures, as can be seen from the high number of genes differentially expressed during this time-point (Figures 3A and 3B). In the short-term response, the number of genes differentially expressed was the highest at 90 min (Figures 3C and 3D). We explored the differentially expressed genes to constrain the genes involved in ES-1 metabolism on different substrates.

**Figure 3.**
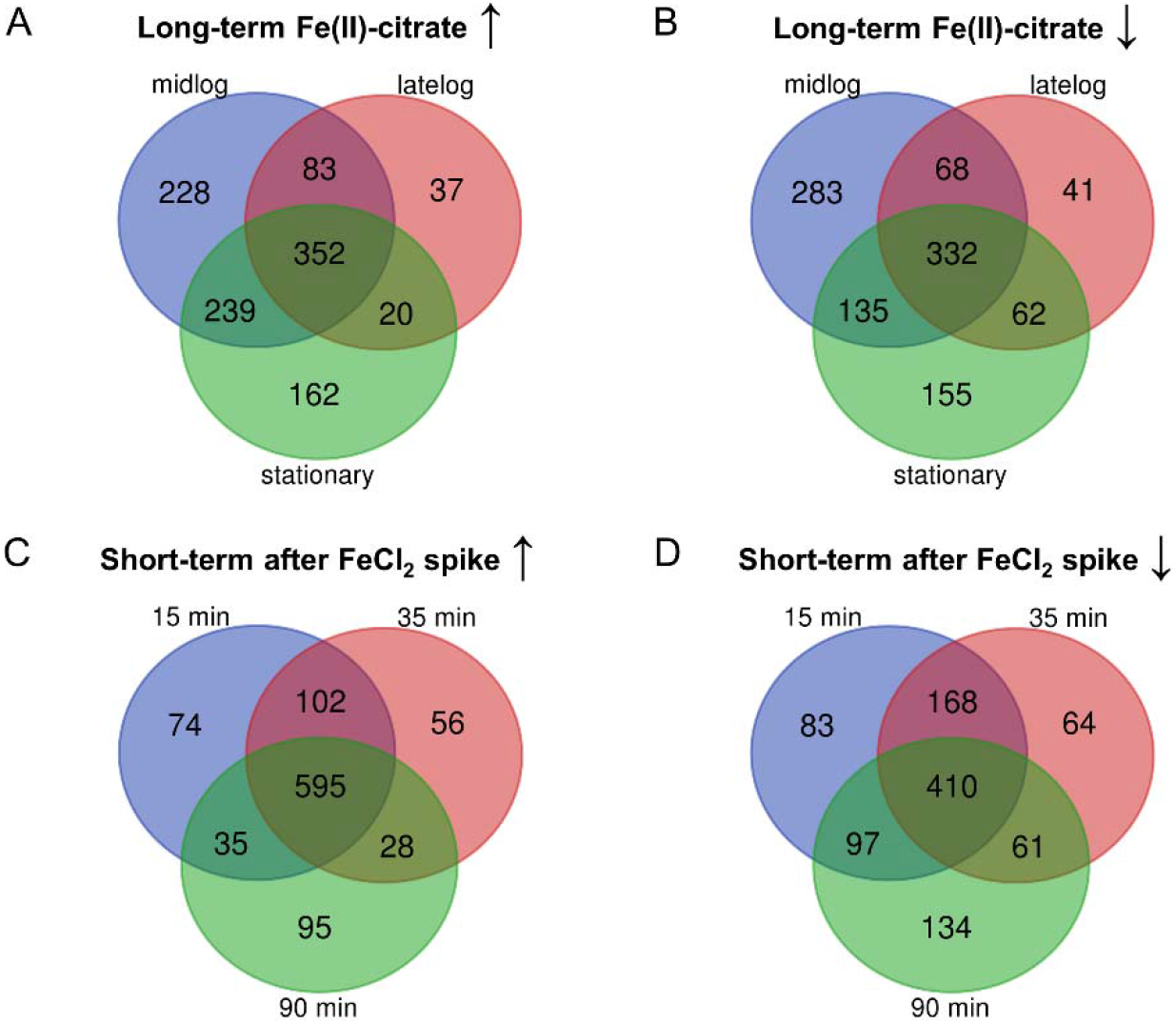
Venn diagram of (A) upregulated genes and (B) downregulated genes in long-term Fe(II)-citrate compared to thiosulfate culture at different growth phases; (C) upregulated genes and (D) downregulated genes at different time points after adding FeCl_2_ compared to time 0 (stationary thiosulfate-grown culture) (p<0.05).

### 2.3 Expression and regulation of thiosulfate oxidases

ES-1 has several thiosulfate oxidation pathways including Tsd (*tsdAB*, Slit_1877-1878), Dsr (*dsrPOJLKMCHFEBA*, Slit_1675-1686) and Sox (*soxBAXYZ*, Slit_1696-1700). Each pathway could produce a different oxidation product, including tetrathionate, elemental sulfur, or sulfate (36). The thiosulfate oxidation product generated by ES-1 is tetrathionate (Figure S1). This suggests the use of *tsdAB* to convert thiosulfate to tetrathionate, since Sox and Dsr pathways are not known to oxidize thiosulfate to tetrathionate (37). However, our transcriptome data showed *tsdAB* genes were neither upregulated nor highly expressed in thiosulfate culture, while *sox* and *dsr* gene clusters were highly expressed and significantly upregulated during thiosulfate growth (Figure 4 and Table S1).

**Figure 4.**
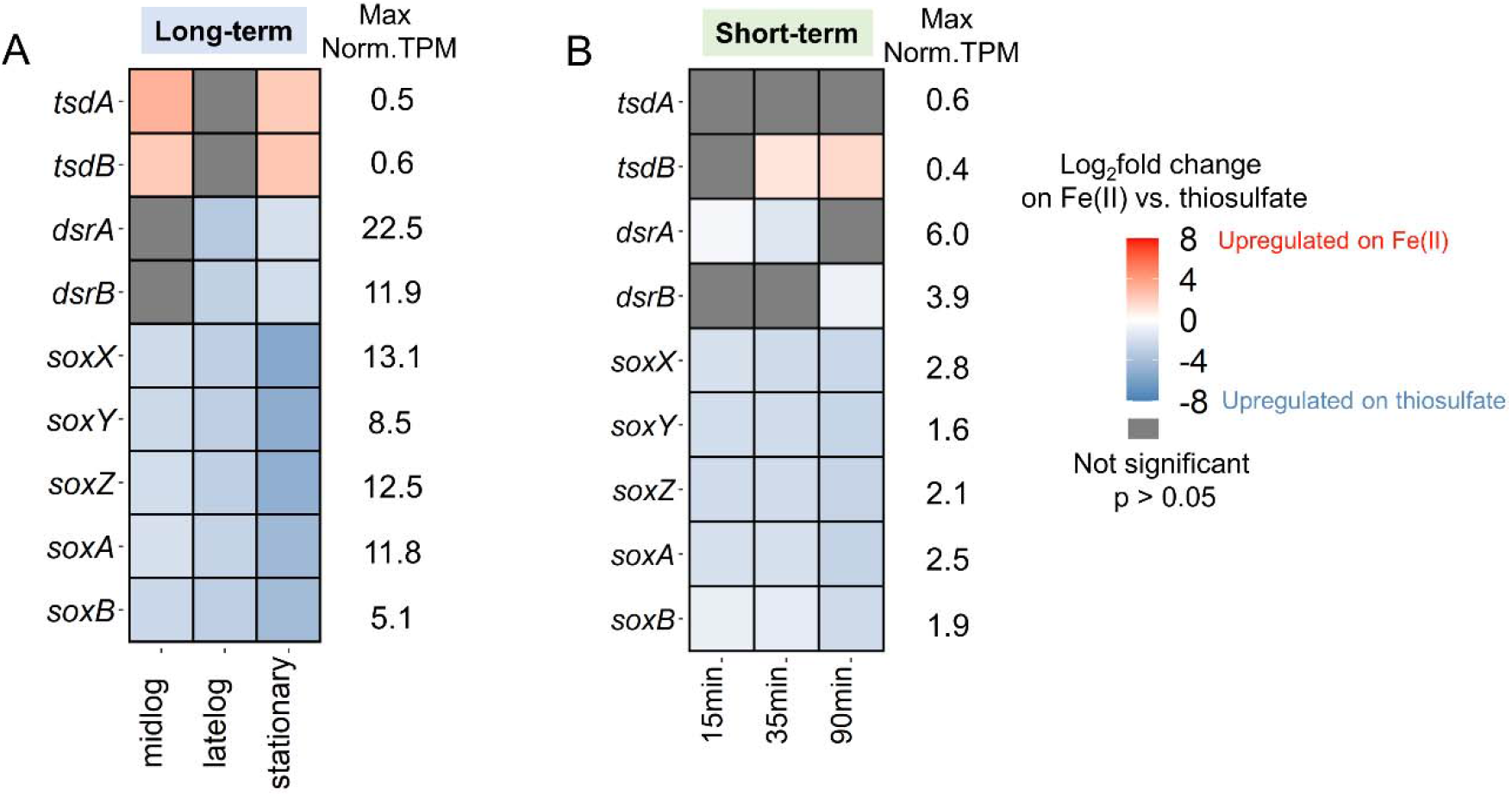
Differential gene expression and maximum constitutive normalized TPM of possible thiosulfate oxidase genes in the (A) long-term and (B) short-term experiments. In the long-term experiment, the reference is thiosulfate-grown ES-1 at different growth phases; in short-term experiment, the reference is Time 0 (stationary thiosulfate grown ES-1). Blue color corresponds to upregulation on thiosulfate.

### 2.4 Expression and regulation of potential Fe(II) oxidases

We examined the possible Fe(II) oxidation genes by transcriptomics and further validated the RNA-Seq results by RT-qPCR. ES-1 has three *cyc2* genes adjacent to one another in the genome separated by 604 and 437 base pair intergenic regions. The three *cyc2* genes are not identical because their amino acid identity varies from 37% to 50%. Each of the four genes (three *cyc2* genes and *mtoA*) behaved differently in response to Fe(II). Overall, the RT-qPCR results compared to the transcriptome showed slightly higher relative expression level than the constitutive normalized TPM values (Tables S2 and S3), possibly because TPM was normalized to multiple genes instead of just *gyrB* in RT-qPCR, but the expression patterns from the two methods were in agreement. In all cultures, *cyc2* expression was generally higher than *mtoA* (Figure 5). In Fe(II)-citrate culture, where the Fe(II)-oxidizing related genes should be expressed, RT-qPCR result shows the three *cyc2* genes expressed on average 359×, 70× and 21× higher than *mtoA* (p < 0.01, Student’s *t*-test) (Figure 5A and Table S2), while the values calculated from constitutive normalized TPM were 1180×, 467× and 157×. Among the three *cyc2* genes, *cyc2_1* showed the highest expression and is one of the top expressed genes in all conditions (Figure 5, Table S3). This result suggests that *cyc2* plays a larger role in Fe(II) oxidation than *mtoA* under the conditions tested.

**Figure 5.**
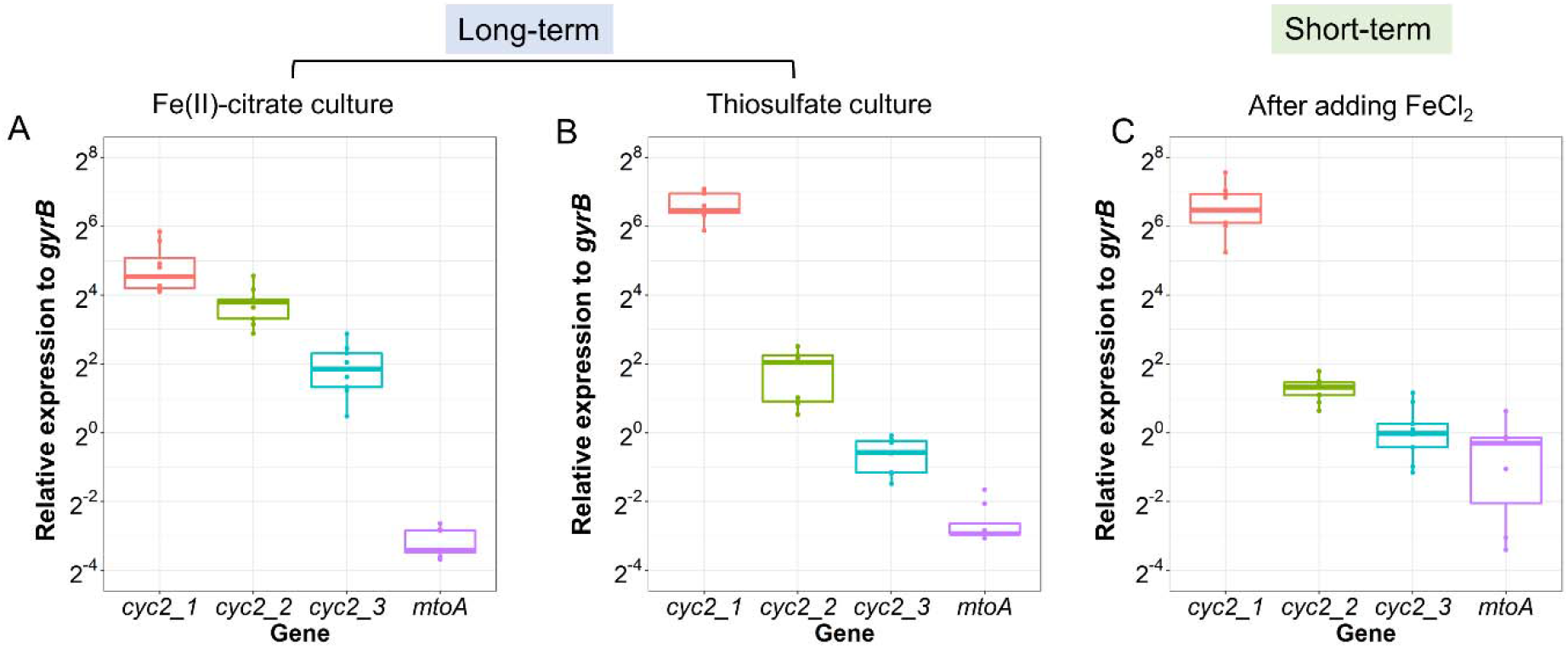
Expression level of the four putative Fe(II) oxidase genes by RT-qPCR in (A) long-term Fe(II)-citrate culture; (B) long-term thiosulfate culture; and (C) short-term after adding FeCl_2_.

The individual responses to Fe(II) varied for each of the three *cyc2* genes. The RT-qPCR results show *cyc2_1* increased in expression level from 106.31 to 145.23 in response to the short-term FeCl_2_ spike (Figure 6, Table S2), though the increase is not significant (p = 0.30). However, the DGE analysis shows a significant upregulation (>1.5 fold, p <0.01) of *cyc2_1* at all three time points after FeCl_2_ addition (Table S3). However, in long-term cultures, *cyc2_1* was downregulated in Fe(II)-citrate cultures compared to thiosulfate cultures. Despite this downregulation, *cyc2_1* remained one of the highest expressed genes in both Fe(II)-citrate and thiosulfate cultures (Table S3). In contrast, the other two *cyc2* copies, *cyc2_2* and *cyc2_3*, were upregulated in long-term Fe(II)-citrate cultures (Figure 6). Moreover, *cyc2_2* was expressed at the same level as *cyc2_1* in Fe(II)-citrate at mid-log phase, when the Fe(II) oxidation activity should be the highest, which suggests *cyc2_2* could play as important a role as *cyc2_1* in Fe(II) oxidation.

**Figure 6.**
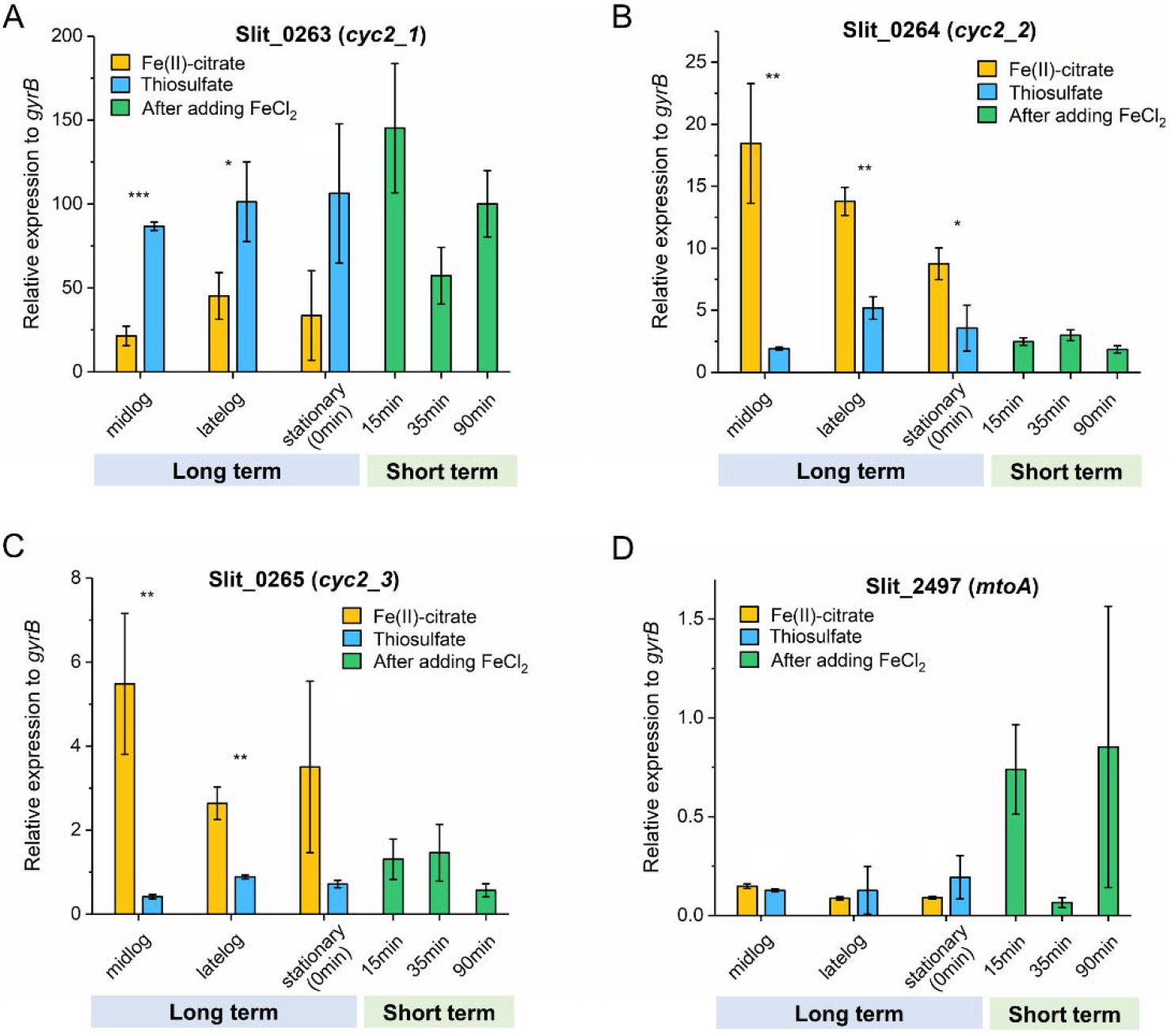
Relative expression by RT-qPCR of three *cyc2* genes and *mtoA* to reference *gyrB* in long-term culturing using Fe(II)-citrate and thiosulfate as substrates and short-term FeCl_2_ spike. In the short-term experiment, time 0 was the stationary phase thiosulfate-grown ES-1. (A) *cyc2_1*; (B) *cyc2_2*; (C) *cyc2_3*; (D) *mtoA*. ^*^ p <0.05, ^**^p <0.01, ^***^p <0.001, unlabeled: not significant from Student’s *t*-test. For the short-term experiment, each time point was compared to 0 min.

### 2.5 Expression of other putative Fe(II) oxidation related genes

In addition to *cyc2* and *mtoA*, we also looked at the expression of other ES-1 genes proposed by He *et al*. (26) as putative Fe(II) oxidases, specifically two “porin-cytochrome c complex” gene clusters (PCC3) which contain a predicted beta-barrel porin, an extracellular multiheme cytochrome (MHC), a periplasmic MHC and a hypothetical protein. ES-1 has two PCC3 gene clusters: Slit_0867-0870 with 17 and 21 heme-binding sites (CXXCH) in the MHC genes Slit_0868 and Slit_0869, respectively, and Slit_1446-1449 with 24 and 28 heme-binding sites in Slit_1447 and Slit_1448, respectively. While the expression level of the second PCC3 gene cluster was higher than the first, both PCC3 gene clusters were downregulated in long-term Fe(II)-citrate grown cells compared to thiosulfate grown cells and in short-term after FeCl_2_ spike compared to time 0 (Table S4). Therefore, neither of the two PCC3 gene clusters appear to be involved in Fe(II) oxidation in ES-1.

We identified a novel gene cluster that was highly expressed (max percentile 97.3) and Fe(II) responsive (Table S4). The gene cluster contains a cytochrome *b* (Slit_1321), a hypothetical extracellular protein (Slit_1322), a monoheme cytochrome class I (Slit_1323), a diheme cytochrome *c* (Slit_1324) and a heat shock protein (Slit_1325). This gene cluster also has homologs in two other neutrophilic Fe(II)-oxidizers, *Gallionella capsiferriformans* ES-2 and *Mariprofundus ferrooxydans* PV-1, consistent with a possible role in Fe(II) oxidation.

Since Cyc2 is an outer membrane Fe(II) oxidase, electrons from Fe(II) need to be transferred to periplasmic cytochromes. We identified two highly expressed monoheme cytochrome class I genes, Slit_1353 and Slit_2042, as candidates for periplasmic electron carriers. Slit_1353 was upregulated in the long-term Fe(II)-citrate culture whereas Slit_2042 was upregulated in short-term after adding FeCl_2_ (Table S4). The *cyc1*_*PV-1*_ homolog Slit_2657 is associated with the *cbb*_*3*_-type terminal oxidase gene cluster and was upregulated in the short-term after FeCl_2_ spike (Table S4).

### 2.6 The expression and regulation patterns in “downhill” electron transfer to terminal oxidases

The electrons obtained from either Fe(II) or thiosulfate may be transferred “downhill” to terminal oxidases (38). In ES-1, the electrons can be passed to a *cbb*_*3*_-type cytochrome oxidase to reduce oxygen (25). There are two *cbb*_*3*_-type cytochrome oxidase gene clusters in the ES-1 genome, which belong to the proximal and distal *cbb*_*3*_ subtrees described by Ducluzeau *et al*. (39) that differ in their subunit composition. The *ccoN* and *ccoO* genes from both *cbb*_*3*_ gene clusters were highly expressed given the high percentile in the gene expression profile, but the constitutive normalized TPM value of distal *cbb*_3_ is ∼3× higher than the proximal *cbb*_*3*_ (Table S5). The DGE analysis shows at mid-log phase, proximal *ccoN* and *ccoO* were upregulated in Fe(II)-citrate grown culture, while the distal *ccoN* and *ccoO* were upregulated in thiosulfate culture (Figure 7). Therefore, each *cbb*_*3*_-type cytochrome oxidase may play a role in accepting electrons from different electron donors.

**Figure 7.**
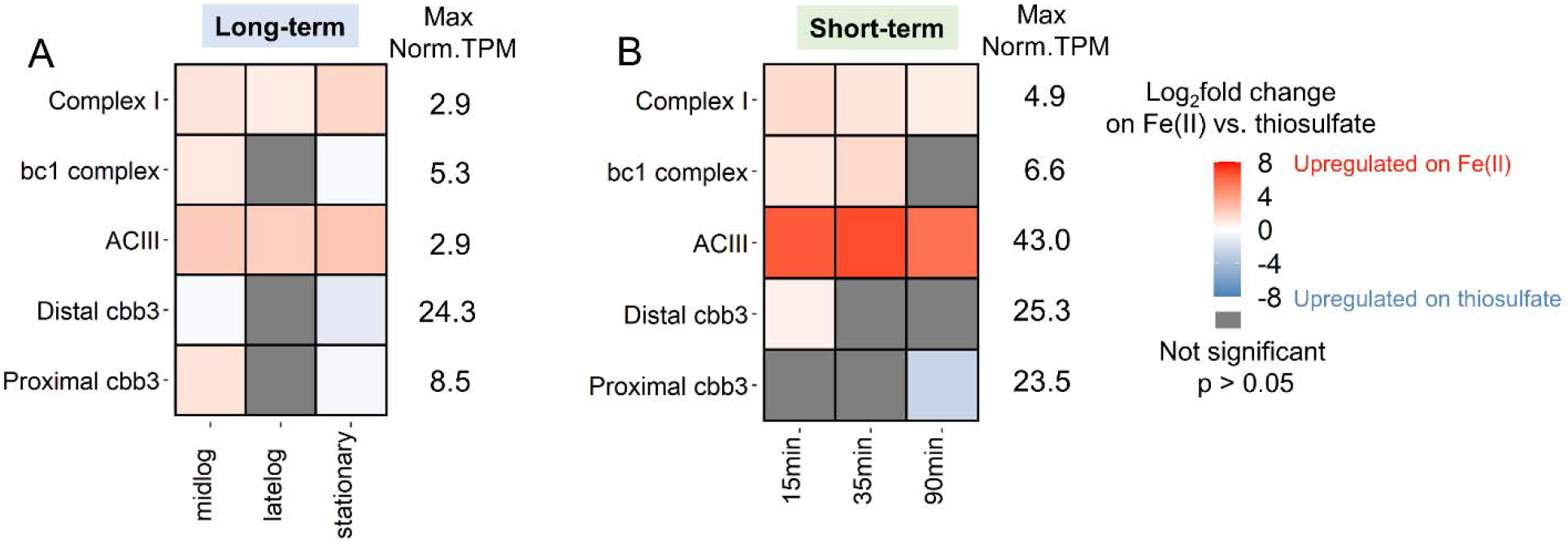
Differential gene expression and maximum values of constitutive normalized TPM of genes involved in “downhill” and “uphill” electron transfer in (A) long-term and (B) short-term experiment. The log_2_fold change in the heatmap is the maximum log_2_fold change in the gene cluster. See Tables S5 and S6 for Log2FC values and Supplemental datasheet for other genes in cluster.

### 2.7 The expression and regulation patterns in “uphill” (reverse) electron transfer

The “uphill” (reverse) electron transport pathway is used to produce reducing equivalents (NADH), which is essential in many biochemical reactions. Importantly, in autotrophic bacteria, NADH participates in CO_2_ fixation to reduce the intermediates, which affects the biomass yield (40). The ES-1 genome encodes both the *bc*_*1*_ complex and Alternative Complex III (ACIII) to potentially carry out the reverse electron transfer to Complex I (40, 41). ACIII genes with high similarity have been found in many other microaerophilic FeOB including *Gallionella capsiferriformans* ES-2 and some *Zetaproteobacteria* FeOB single amplified genomes (SAGs) (13, 21, 25). In our work, the expression level of the *bc*_*1*_ complex (Slit_0130-0132) showed moderate upregulation on Fe(II) (Figure 7, Table S6). DGE analysis showed the ACIII gene cluster was significantly upregulated in Fe(II)-citrate at different growth phases in long-term with the maximum log_2_fold change (log_2_FC) of 2.36 (Table S6). After adding FeCl_2_ to the thiosulfate culture, the ACIII gene cluster showed a sharp upregulation with the maximum log_2_FC of 6.84 (Table S6). Meanwhile, the gene cluster of NADH dehydrogenase (Complex I) was also upregulated in long- and short-term response to Fe(II) substrates (Figure 7). The upregulation of “uphill” electron transfer chain could result in more production of NADH during Fe(II) oxidation, to be further used in various biochemical reactions.

### 2.8 Regulation patterns of CO_2_ fixation

CO_2_ fixation is an important NADH sink (42, 43). In ES-1, CO_2_ fixation is achieved through Calvin Benson (CBB) cycle, in which RuBisCO proteins are the key enzymes (25). ES-1 encodes two types of RuBisCO: Form I *cbbLS* (Slit_0985-0986) and Form II *cbbMQ* (Slit_0022-0023). The Form II enzymes have a higher affinity for O_2_ compared to Form I, which allows CO_2_ fixation at lower O_2_ partial pressures (44). The maximum normalized TPM of *cbbMQ* was over 100× higher than the *cbbLS*, which indicates *cbbMQ* might play a major role in ES-1 CO_2_ fixation. RuBisCO (*cbbMQ*) and most of the other genes involved in the CBB cycle were upregulated in long- and short-term response to Fe(II) (Figure 8). Moreover, the short-term response showed a sharp upregulation of *cbbMQ*, with a log_2_FC greater than 5, which is consistent with the ACIII regulation pattern.

**Figure 8.**
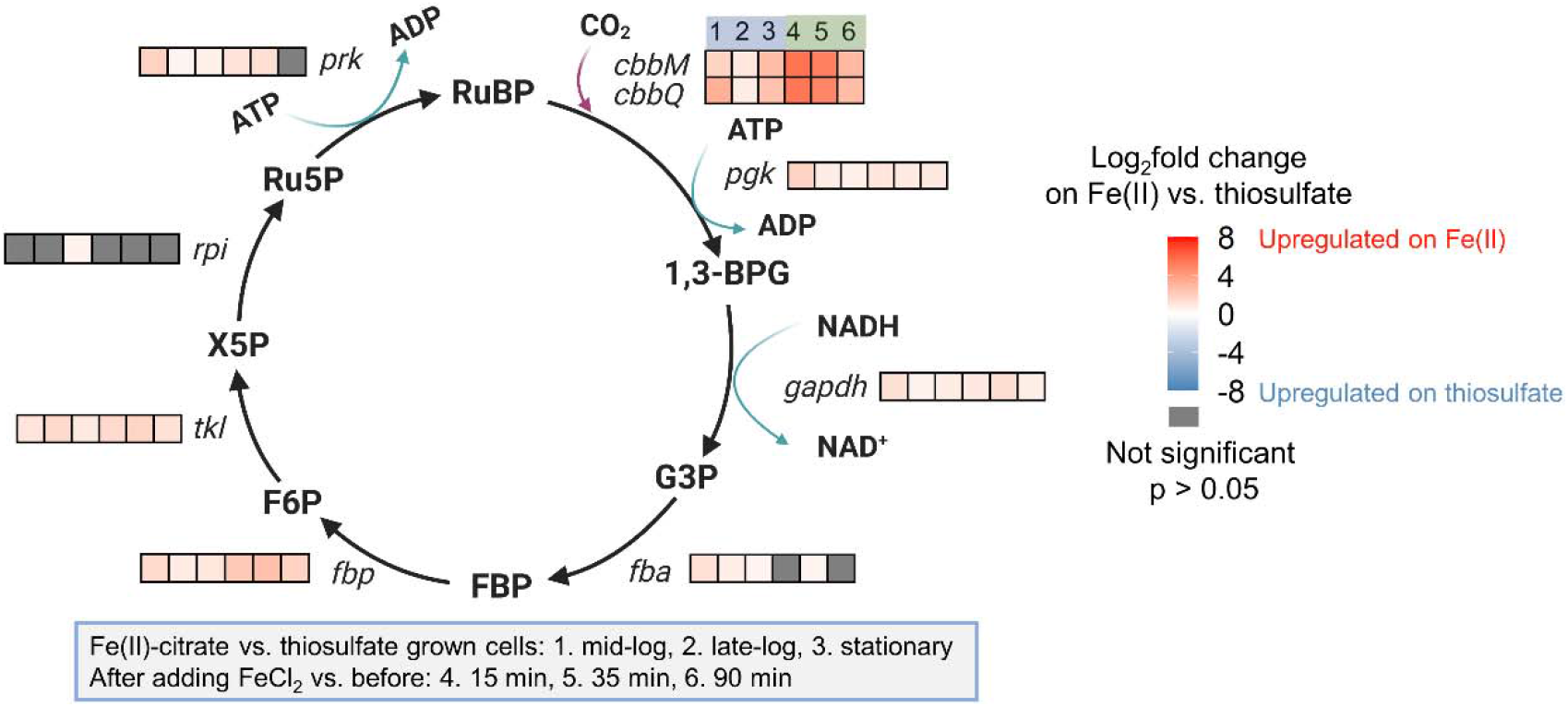
Differential gene expression of the genes involved in CBB cycle. RuBP (Ribulose 1,5-bisphosphate), 1,3 BPG (1,3-bisphosphoglycerate), G3P (Glyceraldehyde 3-phosphate), FBP (Fructose 1,6-bisphosphate), F6P (Fructose 6-phosphate), X5P (Xylulose 5-phosphate), Ru5P (Ribulose 5-phosphate).

### 2.9 Phage genes

ES-1 has two clusters of prophage genes, and these genes are among the most responsive to the change in substrates. The first phage cluster (Slit_0188-Slit_0249; 39.7 kb) belongs to a Mu-like prophage family, and a subset of these genes have homology to a prophage in the marine microaerophilic FeOB, *Mariprofundus ferrooxydans* PV-1 (25). The second phage gene cluster (Slit_1888-Slit_1969; 54.3 kb) does not have homologs in PV-1 and the majority of the genes have hypothetical or unknown function. The DGE analysis showed the first phage cluster is Fe(II)-responsive (upregulated in Fe(II)-citrate culture) with the maximum log_2_FC of 7.90, whereas the second phage cluster is thiosulfate-responsive (upregulated in thiosulfate culture). These data do not illuminate whether these gene expression trends are due to induction of the prophage within some fraction of the ES-1 population into an active lytic cycle, or whether they are the result of lysogenic conversion (i.e. host expression of integrated prophage genes). However the result does suggest that the phage gene clusters may represent auxiliary metabolic genes (45) that could play a role in ES-1 Fe(II) and thiosulfate oxidation, and/or that the phage activation itself may be responsive to the availability and oxidative metabolism of these compounds.

## 3. Discussion

*Sideroxydans lithotrophicus* ES-1 was isolated as an Fe(II)-oxidizer and was later discovered to also grow by thiosulfate oxidation (25, 46). This make it unusual amongst the *Gallionellaceae*, in which most other isolates are obligate Fe(II)-oxidizers (*Gallionella, Ferriphaselus*, and *Ferrigenium* spp.). Previous metagenomic studies have shown *Sideroxydans* are abundant in environments with both Fe(II) and reduced S species (21–23, 47). Our data shows ES-1 has higher yields on thiosulfate (Figure 1A and Figure S2) and can grow simultaneously on Fe(II) and thiosulfate (Figures 1A, 1B and 1C). Our growth studies show that the addition of thiosulfate can boost *Sideroxydans* growth in the presence or absence of Fe(II), giving it an advantage over obligate FeOB in niches where both Fe and S are available.

Because different pathways are used to oxidize Fe(II) vs. thiosulfate, cells might conserve energy by expressing only the genes necessary for the available or desired electron donor(s). We performed experiments in which we switched the substrate and observed a lag in substrate consumption when switching from Fe(II) to thiosulfate (Figure 1C), but no lag when switching from thiosulfate to Fe(II) (Figure 1D). This is consistent with the expression patterns of Fe(II)- and thiosulfate-oxidizing genes. There was relatively high expression of Fe(II)-oxidizing gene *cyc2_1* during the thiosulfate-only growth (Figures 5 and 6), whereas S oxidation genes were expressed at low levels during Fe(II) oxidation and significantly upregulated on thiosulfate (Figure 4). This contrast suggests that *Sideroxydans* maintains readiness to oxidize Fe(II), while thiosulfate oxidation serves as a secondary, supporting metabolism either in the presence or absence of Fe(II). In this case, despite the ability to oxidize a sulfur compound, the *Sideroxydans* niche appears to be primarily in Fe(II) oxidation, consistent with its known environmental distribution.

The differential gene expression on Fe(II) vs. thiosulfate gives us insight into the genes that may be specific to each metabolism. ES-1 upregulated *dsr* and *sox* genes on thiosulfate, despite the expectation that *tsd* would be upregulated, since thiosulfate is incompletely oxidized to tetrathionate. The Dsr and Sox pathways are not known to produce tetrathionate, and ES-1 has not yet been shown to oxidize sulfide, so the specific roles of the ES-1 *dsr* and *sox* genes require further investigation, and might prove to have different functions in ES-1 compared to other sulfur-oxidizing microbes. Still, the *dsr* and *sox* gene expression does correspond to thiosulfate oxidation activity in ES-1.

The ES-1 genome encodes several proposed Fe(II) oxidation pathways including those involving Cyc2, Mto, and PCC3. Overall, the *cyc2* expression levels were much higher than *mtoA* or PCC3 in the presence of Fe(II) and *cyc2_1* was among the top expressed genes (>99 percentile; Tables S3 and S4), suggesting the importance of *cyc2* in Fe(II) oxidation. The presence of three *cyc2* genes further suggests the utility since multiple copies might enable ES-1 to increase its Fe(II) oxidation capacity or efficiency (48–50). The three *cyc2* genes had different responses to Fe(II) in the short- and long-term experiments, suggesting that each gene plays a distinct role. The first copy, *cyc2_1*, increased expression on Fe(II) in the short-term experiment, but overall was highly expressed under all conditions, including the thiosulfate grown culture. This continuous expression may imply regulation at the protein expression level. However, thiosulfate-grown cells were able to immediately oxidize Fe(II) (Figure 1D), suggesting that Cyc2 is indeed expressed, possibly to maintain “readiness” and may also act as an Fe(II) sensor. Addition of Fe(II) triggers the expression of *cyc2_2* and *cyc2_3* (Figure 6, Tables S2 and S3), which would allow for more efficient transcription and translation when Fe(II) levels are high. In all, the *cyc2* expression levels and patterns support the model of a Cyc2-based Fe(II) oxidation pathway.

For neutrophilic FeOB like ES-1, the Cyc2 Fe(II) oxidation pathway was originally deduced from comparative genomics, and recent work has verified the Fe(II)-oxidizing function of Cyc2_PV-1_ (29), a relatively close homolog of Cyc2_ES-1_. Apart from Cyc2, the rest of the pathway has not been tested in neutrophilic FeOB isolates (25). Here, we constrained and updated the model based on gene expression levels and differential expression (Figure 9). We identified likely periplasmic cytochromes that could act as electron carriers to transfer electrons from an outer membrane Fe(II) oxidase to the “uphill” (Slit_1353, Slit_2042) and “downhill” (Slit_2657) electron transfer pathways (Figure 9, Table S4). Both long-term and short-term data showed upregulation of RET and CO_2_ fixation related genes in the presence of Fe(II) (Figures 7 and 8), which suggests electrons from Fe(II) tend to pass through RET to CO_2_. Notably, genes for ACIII and RuBisCO were massively upregulated after the Fe(II) pulse. ACIII gene clusters in Fe(II)-oxidizing *Betaproteobacteria* and *Zetaproteobacteria* have a high degree of similarity, leading to a hypothesized role in Fe(II) oxidation, though the actual function remained unclear (26, 51–53). Our result suggests ACIII mainly participates in RET to provide reducing equivalents for the CO_2_ fixation. In this model, CO_2_ is an important electron sink to maintain redox balance, similar to photoautotrophic FeOB (42, 43).

**Figure 9.**
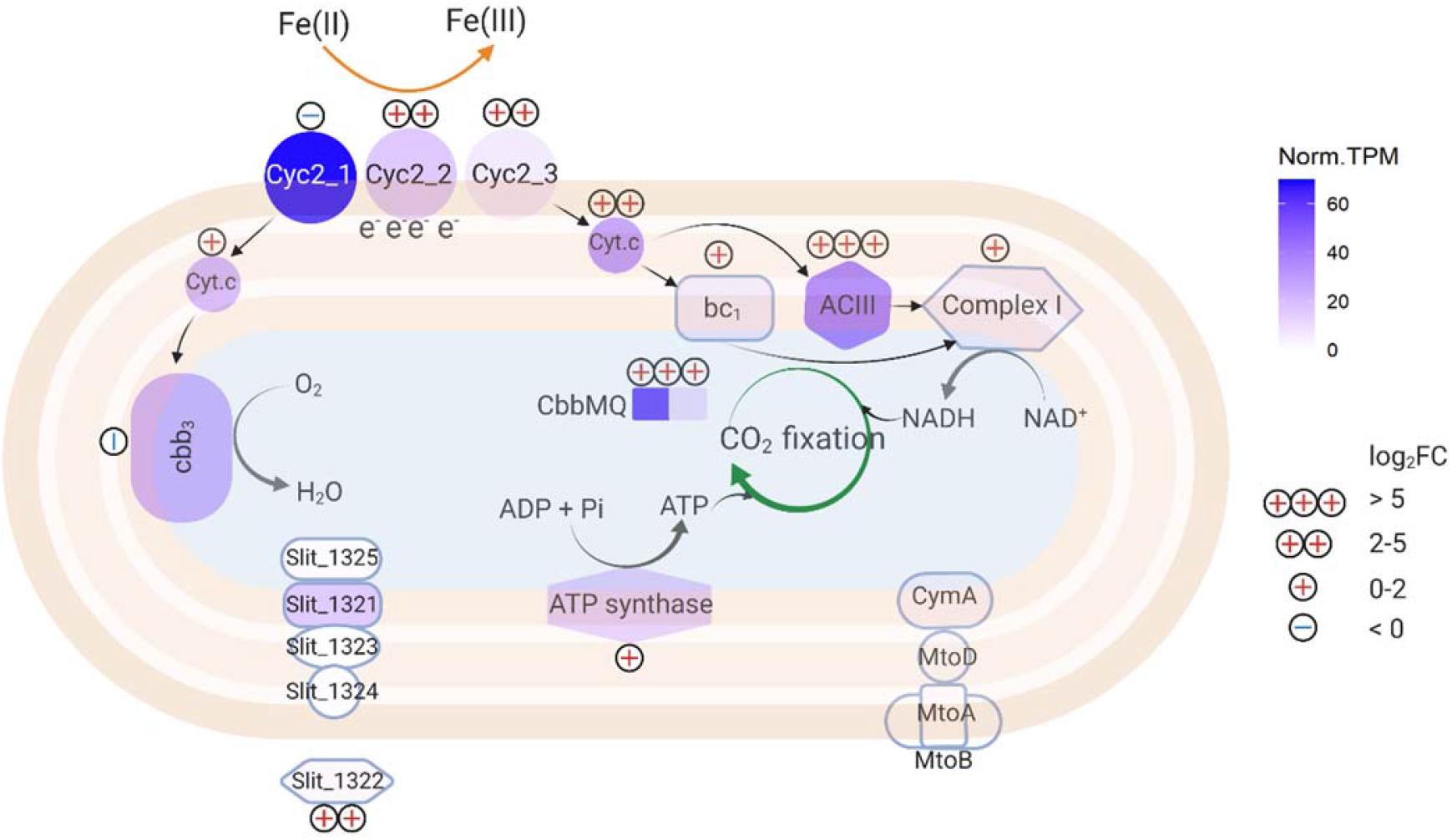
Updated Fe(II) oxidation pathway in ES-1. Norm.TPM represents the maximum normalized TPM using constitutively expressed genes; log_2_FC represents the maximum log_2_ fold change in Fe(II) culture as compared to thiosulfate culture (included both long-term and short-term data). The protein location of Slit_1321-1325 was predicted using PSORTb version 3.0.2 (54). The figure was created using Biorender.com.

In all, our results provide a clearer view of *Sideroxydans lithotrophicus* ES-1’s physiology and gene responses to Fe(II) and thiosulfate oxidation, which may be applied to understanding environmental Fe(II) oxidation. Detecting microbial/biotic Fe(II) oxidation in the environment has historically been difficult since biotically- and abiotically-formed Fe(III) oxides are not necessarily distinct by either minerology or isotope composition (55). There is now the potential to use the expression of Fe(II) oxidation genes to signal environmental Fe(II) oxidation if gene expression can be linked to activity. The most obvious candidates for Fe(II) oxidation gene markers are Fe(II) oxidase genes. In the case of ES-1, this is *cyc2*, and in fact, *cyc2* homologs are common across the Fe(II)-oxidizing *Gallionellaceae* and *Zetaproteobacteria* (26, 51, 53, 56), and Cyc2 from multiple organisms has been shown to oxidize Fe(II) (27, 29, 57). The *cyc2* genes are highly expressed in Fe(II)-oxidizing environments, as shown by metatranscriptomic studies on marine *Zetaproteobacteria* iron mats (53) and an Fe-rich aquifer dominated by *Gallionellaceae* (of which *Sideroxydans lithotrophicus* ES-1 is a member) (7). Although microbial activity and gene expression can be equated with Fe(II) oxidation activity in obligate FeOB like many *Zetaproteobacteria*, our work revealed a more complicated picture in a facultative FeOB, which also expressed *cyc2* under non-Fe(II)-oxidizing conditions. In environments with facultative FeOB, *cyc2* or another Fe(II) oxidase gene may not be sufficient for monitoring Fe(II) oxidation activity. Instead, we can monitor gene expression for the full Fe(II) oxidation pathway, including forward and reverse electron transport (Figure 9). Our work suggests that the full pathway can be used as a multi-gene indicator of Fe(II) oxidation activity, and coupled monitoring of C fixation genes can further signal Fe(II)-oxidation driven C fixation, thus showing interconnected Fe-C biogeochemical cycling in environment.

## 4. Materials and Methods

### 4.1 Cell cultivation

*Sideroxydans lithotrophicus* ES-1 was grown in Modified Wolfe’s Minimal Medium (MWMM), which contains 1 g/L NH_4_Cl, 0.5 g/L MgSO_4_•7H_2_O, 0.2 g/L CaCl_2_ and 0.05 g/L K_2_HPO_4_. The MWMM was buffered with 20 mM 2-(N-morpholino)ethanesulfonic acid (MES) and adjusted to pH 6.0. The liquid media was deoxygenated by purging with nitrogen gas. Media was supplemented with Wolfe’s vitamin and trace mineral solution (ATCC, 2672) in a 1:1000 ratio after autoclaving (58). The headspace was maintained at 2% O_2_, 20% CO_2_ and 78% N_2_ by flushing daily. Culturing on Fe(II)-citrate was sustained with a one-time addition of sodium citrate to a final media concentration of 5 mM and a daily addition of FeCl_2_ stock solution. Cell cultivation on thiosulfate was performed by a one-time addition of sodium thiosulfate. Substrate concentration was varied (Fe: 100-750 μM/day, thiosulfate: 0.5-10 mM) for optimizing growth (Figure S2). The cell number was determined by counting Syto13-stained cells under fluorescent microscopy using a Hausser counting chamber.

### 4.2 ES-1 substrate consumption experiment

Ferrous chloride (FeCl_2_)-grown cells were inoculated into MWMM media with both FeCl_2_ and thiosulfate, FeCl_2_ only, or thiosulfate only. The uninoculated media with both FeCl_2_ and thiosulfate was used as the abiotic control. All treatments were performed in triplicate. Substrates (FeCl_2_ and sodium thiosulfate) were amended daily to maintain a concentration of 500 μM. The headspace was flushed with gas mix (O_2_:CO_2_:N_2_=2:20:78) every day. Cell number was recorded every two days by direct cell counting. Fe(II) and thiosulfate concentration were determined every day before and after the substrate supplement. The biotic Fe(II) and thiosulfate consumption per day was calculated by subtracting the substrate loss in the uninoculated control from the substrate loss in the live culture.

### 4.3 Chemical analysis

The Fe(II) concentration was determined by spectrophotometric ferrozine assay (59, 60). To ensure oxygen was maintained between 20-30 µM, the oxygen concentration in the cultures was measured using a Firesting oxygen meter coupled with a fiber optic probe, and a sensor spot inside the serum bottle (Pyro Science). Thiosulfate concentration was determined by mixing samples with the same volume of 1 mM iodine solution, then the absorbance at 350 nm was measured (61). The sulfur speciation was identified using high performance liquid chromatography (HPLC). Samples were separated on an Alltech Organic Acids 5 µm column (150 × 4.6 mm) eluted with 25 mM potassium phosphate buffer at 1 ml/min, pH 2.5, on a Shimadzu Class-10VP HPLC. Detection was at 210 nm. Thiosulfate and tetrathionate standards were prepared and diluted in growth medium to generate a standard curve and calibrate the retention time for each compound.

### 4.4 Transcriptome experiment culturing and sampling

Transcriptome analysis was used to investigate the genes expressed during Fe(II) and thiosulfate oxidation. To obtain sufficient biomass for RNA extraction, the culture was scaled up to 250 mL of media in 500 mL glass culture bottles. We investigated both long-term and short-term responses of ES-1 to different Fe(II) substrates. To investigate the long-term response, we cultivated ES-1 using Fe(II)-citrate or thiosulfate as described above, then sampled different growth phases. Fe(II)-citrate grown cells were sampled on day 2 (mid-log), day 5 (late-log) and day 7 (stationary). Samples from thiosulfate-grown cells were taken on day 3 (mid-log), day 7 (late-log) and day 10 (stationary). Since the Fe(II)-citrate grown cultures had lower cell density, three culture bottles were combined (750 mL total) for each sample. The thiosulfate-grown cultures had higher cell density, therefore, only one bottle (250 mL) of culture was harvested for each sample. To investigate short-term, rapid gene expression responses to Fe(II), we spiked FeCl_2_ stock solution into stationary phase thiosulfate-grown ES-1 culture to a final concentration of 100 μM and the killed-cell control was created by adding sodium azide into the culture to a final concentration of 5 mM. Samples were taken at 0, 15, 35, 60 and 90 min for ferrozine assay to track the Fe(II) oxidation kinetics. Meanwhile, culture bottles were sacrificed for RNA extraction at the beginning (15 min), middle (35 min) and end of the Fe(II) oxidation (90 min). For each time point in the long-term and short-term experiment, samples were collected in biological triplicate. To protect cellular RNA from degradation, 1/10 volume of stop solution (buffer-saturated phenol : absolute ethanol = 1: 9, v/v) was added to the culture (62) and cells were harvested by collecting the culture on a 0.22-micron membrane (Millipore GTTP). The filter membranes with cells were then cut into small pieces and stored at -80 °C until RNA extraction.

### 4.5 RNA extraction

The cells were lysed by high speed vortexing for 2 minutes using lysing matrix E tubes (MP Biomedical) in buffer RLT (Qiagen) amended with 1% 2-mercaptoethanol. The lysate was centrifuged at 4000 × *g* for 10 min in 4 °C. The supernatant was collected and extracted using the Qiagen Micro RNeasy kit following the instructions provided by the manufacturer. The samples were then treated with the Turbo Dnase kit (Invitrogen), to remove genomic DNA. The ribosomal RNA in the total RNA samples was depleted using MICROBExpress Bacterial mRNA purification kit (Invitrogen). The rRNA depleted samples were further purified and concentrated using Zymo RNA Clean and Concentrator kit. The RNA concentration was quantified on a Qubit Fluorometer using the Qubit RNA HS assay kit (Invitrogen). The RNA quality was determined on the Agilent Fragment Analyzer at the University of Delaware DNA Sequencing & Genotyping Center.

### 4.6 Reverse transcription quantitative PCR (RT-qPCR)

Total RNA (20 ng) was used as template to synthesize cDNA using Maxima First Strand cDNA Synthesis Kit (Thermo Scientific). Gene expression of three *cyc2* genes (Slit_0263, Slit_0264, Slit_0265) and *mtoA* (Slit_2497) was quantified using RT-qPCR with *gyrB* (Slit_0003) as a reference gene. Primers for the target genes were designed (sequences in Supplementary Table S7). Quantabio SYBR Green supermix and Bio-Rad CFX96 Real-Time PCR System were used to perform the quantification assays. Standard curves were generated using cloned plasmids containing the target DNA fragments with the concentration range from 500 ag/μL to 50 pg/μL. The target genes were amplified using the following program: 95°C 5 min, followed by 40 cycles of 95°C 15 s, then the indicated annealing temperature of each primer set for 45 s (Table S7), 72°C for 30 s and followed by a melt curve analysis from 55-65°C. The gene copy numbers were calculated using the quantification cycle (C_q_) values and the standard curves for each gene. Each biological replicate was run in two technical replicates.

### 4.7 RNA sequencing and analysis

The library preparation and sequencing were performed at the University of Delaware DNA Sequencing & Genotyping Center. The NEXTFLEX Rapid Directional RNA-Seq Kit was used for library preparation. Sequencing was performed on Illumina NextSeq 550 Sequencing system with a read length of 1×76 bp. The transcriptomic reads were quality controlled using FastQC v0.11.9 (Babraham Bioinformatics). The low-quality reads were trimmed using Trim Galore! v0.6.4 with a cutoff Q score of 28. After trimming, the average reads per sample was 19.6 M. Then the trimmed reads were mapped to the ES-1 genome (GenBank accession No. NC_013959) using bowtie2 v2.3.5.1 (63). The overall average alignment rate among all the samples was 98.3%. Next, samtools v1.9 was used to sort and index the aligned file (64). The sorted .bam files were counted using Htseq-count (65) with annotated ES-1 gff file. To evaluate the gene expression level, we first normalized the read counts to transcripts per million (TPM), which accounts for both gene length and library size, then TPM values were further normalized by dividing by the average TPM values of six constitutively expressed genes (*gyrB, gyrA, adk, rho, era, gmk*) (35). The percentile of gene expression was calculated based on the constitutive normalized TPM. The output counting file was used to analyze the differential gene expression using Bioconductor DESeq2 v1.26.0 (66), and p-values were FDR-adjusted using the Benjamini-Hochberg (BH) correction (67). For the long-term experiment, simple pairwise comparison was performed between Fe(II)-citrate and thiosulfate grown cells at each different growth phase with thiosulfate grown cells as the control. For the short-term experiment, the control was time 0 (thiosulfate-grown cells in stationary phase), and other time points after Fe(II) addition were pairwise compared with time 0. Genes with zero counts were removed from the DGE analysis. Genes with p-values <0.05 were considered differentially expressed. Heatmaps were made in R using ggplot2 (68). A master spreadsheet with raw counts, constitutive normalized TPM values and DGE analysis results is provided as a supplemental file. Raw RNA-Seq reads were uploaded to NCBI BioProject GEOarchive with the accession number GSE181717.

## Acknowledgements

This work is funded by NASA Exobiology (80NSSC18K1292 to CSC), NSF Geobiology and Low Temperature Geochemistry (EAR-1833525 to CSC and SWP), ONR grant N00014-17-1-2640 (to CSC), and the Joanne Daiber Fellowship (to NZ). We would like to acknowledge the University of Delaware DNA Sequencing Center for RNA sequencing and Delaware INBRE for transcriptomic data processing using Biomix (NIH P20 GM103446). We thank Thomas Hanson for his help in characterizing thiosulfate oxidation products and David Emerson for his comments on an early version of this manuscript.

